# The conversion of systolic volume into systolic pressure (the development of numerical model)

**DOI:** 10.1101/011536

**Authors:** Yuri Kamnev

## Abstract

The direct proportional trend of the transition from systolic volume to systolic pressure can be proved theoretically with the help of circulatory numerical model. The model permits to link calculable end-diastolic volume with the observable interrelations between systolic and diastolic pressures. The outstripping growth of systolic pressure (when diastolic pressure is rising with steady increment) is deduced and the analogous phenomenon can be observed at clinical gradation of arterial hypertension.

## Introduction

The present article ought not to be considered the independent theoretical investigation because it just branches out from the previous, basic article and develops the only one section of the model, hence, the references to [1] will imply continually. This section needs the detailed description and the special proof but it would overload the main demonstration of the basic article [1] and we decided to represent the section separately.

## Methods

At present simulation we have equalized the end-diastolic volume (which is a calculable value by means of model presented at the article [1]) and systolic pressure; more strictly, we have supposed direct proportionality (with coefficients of proportionality equal to1) between the end-diastolic volume, L, systolic volume, V_syst_, and systolic pressure, P_syst_, and we are in need to prove it, i.e., to prove the permissibility of such substitution. In other words, the process described at the model (where the above substitution works) must correspond to some physiological phenomenon. Practically it means the following, firstly: the deviating parameters that reveal the operation of the model (elevation or falling of the resistance, the change of the value of normal rhythm, acceleration of the volume flow rate, which all can not be observed easily) must be associated with fluctuations of diastolic pressure at different modes of circulation (norm-, tachy- or brady-modes) since the compound parameter *Q · ΣR* (where Q is the volume flow rate and ΣR is the resistance) can be interpreted as diastolic pressure, P_diast_, which is quite available for measuring in practice. Secondly, the calculated parameter L (the end-diastolic volume, which is also hardly observable) must behave identically to systolic pressure, P_syst_,which can be easily measured in pair with P_diast_, i.e., almost simultaneously. Physiological parameter that combines the resistance (together with volume flow rate, i.e., *Q* · *Σ R*) and systolic pressure is the ratio called “arterial pressure” 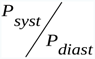, where P_diast_ denotes the variable resistance ΣR (which we use in calculations) multiplied by constant value of Q. Variations of ΣR coincide with changes of P_diast_ in case when the volume flow rate is equal to 1; in case when Q differs from 1 it operates as constant factor that shifts the values of P_diast_ proportionately. The derivative parameter is a pulse pressure (the difference between systolic and diastolic pressures) which is convenient due to its expression by one number and its independence from certain values of systolic and diastolic pressures. Therefore, the task of simulation is to trace the sequence of values of arterial pressure as a response to series of values of diastolic pressure deviated with steady increment or decrement; the achieved ratios of arterial pressure, supplemented by series of pulse pressure, must be assessed in accordance with degrees of arterial hypertension observed in clinical practice and sports medicine.

We ought to remind how we interpret the measuring of systolic and diastolic pressure at the same anatomical region, i.e., at the region of initial arteries which are located between the ventricle and resistive arteries. The approach is to frame the triple parameter where the pressure is linked to the certain anatomical portion of pipe and to the phase of cardiac cycle [1]. Respectively, the systolic pressure is considered essential to the initial arteries but registration of diastolic pressure at this initial portion of pipe relates to the effect of conduction of pressure from the region of resistive arteries where diastolic pressure is originated.

## Results

Tab.1 is based on data from Fig.1 [1]; we have used the data that regard to the steps of reorganization of norm-mode of circulation (t_N_ =6, P_N_ =6, Q=1.0) as a response upon the rising and falling of the resistance but excludes the restoration of each step to the previous one. Three steps of response of L (which is assumed numerically equal to P_syst_) upon the rising of the resistance from the point of normal value R_N_ = 30 (increment ΔR = 10) and two steps of response of L upon the falling of the resistance (decrement ΔR = -10) are represented. Tab.1 shows the series of ΣR which are numerically identical with the values of P_diast_ shown either in the line “Resistance” or in the line “Arterial pressure”, and the series of L from Fig.1[1] is numerically identical with the values of P_syst_ which are shown in the line “Arterial pressure”. It is evident that pulse pressure is not a fixed value when the resistance is changing; pulse pressure grows linearly with increment of growth equal to 2.

**Table 1.**
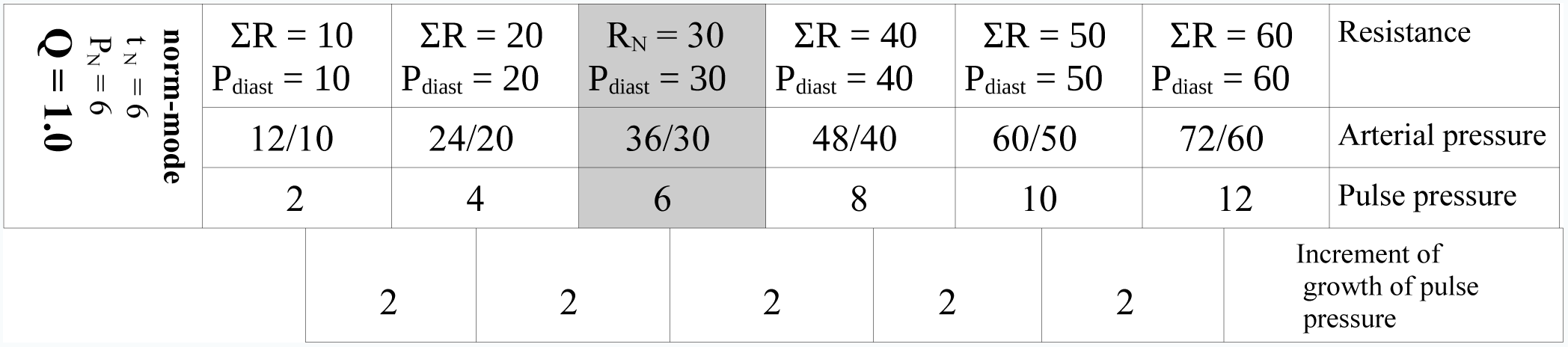
The steady rising of the resistance (which is identical to diastolic pressure when Q = 1.0) results in growth of pulse pressure; the represented norm-mode of circulation (which basic normal rhythm is not tachy-rhythm or brady-rhythm) is shortly described by information in the upper left cell; the gray column indicates the state of circulation when it is not deviated by shift of the resistance.

Tab.2 is based on data from Fig.2 [1]; we have used the data that regard to the steps of reorganization of tachy-mode of circulation (t_N_ =3, P_N_ =15, Q=1.0) as a response upon the rising and falling of the resistance but excludes the restoration of each step to the previous one. Three steps of response of L (which is assumed numerically equal to P_syst_) upon the rising of the resistance from the point of normal value R_N_ = 30 (increment ΔR = 10) and two steps of response of L upon the falling of the resistance (decrement ΔR = -10) are represented. Tab.2 shows the series of ΣR which are numerically identical with the values of P_diast_ shown either in the line “Resistance” or in the line “Arterial pressure”, and the series of L from Fig.2 [1] is numerically identical with the values of P_syst_ which are shown in the line “Arterial pressure”. It is evident that pulse pressure is not a fixed value when the resistance is changing; pulse pressure grows linearly with increment of growth equal to 5.

**Table 2.**
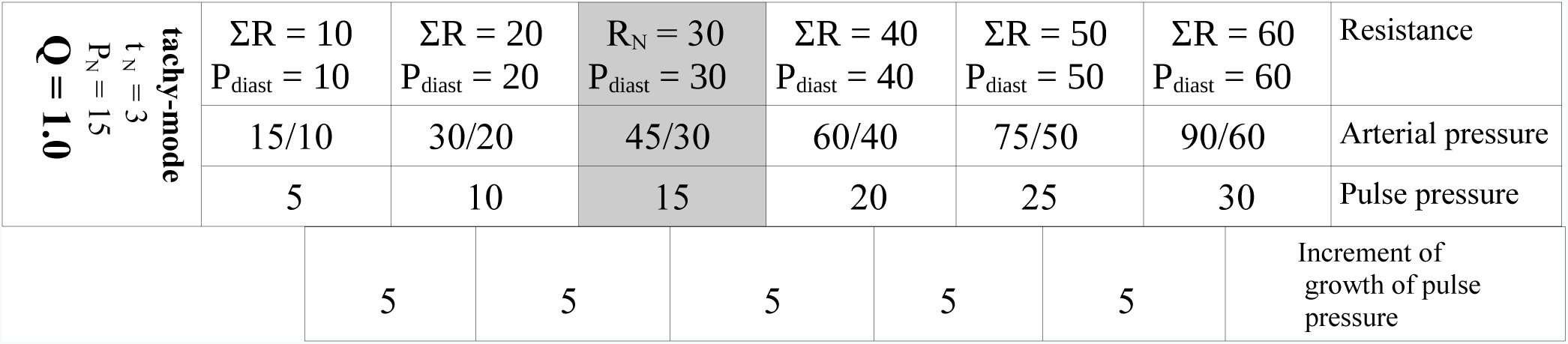
The steady rising of the resistance (which is identical to diastolic pressure when Q = 1.0) results in growth of pulse pressure; the represented tachy-mode of circulation is shortly described by information in the upper left cell; the gray column indicates the state of circulation when tachy-mode is not deviated by shift of the resistance (it is the starting point, i.e., the normal state for tachy-mode).

Tab.3 is based on data from Fig.3 [1]; we have used the data that regard to the steps of reorganization of brady-mode of circulation (t_N_ =9, P_N_ =3.75, Q=1.0) as a response upon the rising and falling of the resistance but excludes the restoration of each step to the previous one. Three steps of response of L (which is assumed numerically equal to P_syst_) upon the rising of the resistance from the point of normal value R_N_ = 30 (increment ΔR = 10) and two steps of response of L upon the falling of the resistance (decrement ΔR = -10) are represented. Tab.3 shows the series of ΣR which are numerically identical with the values of P_diast_ shown either in the line “Resistance” or in the line “Arterial pressure”, and the series of L from Fig.3[1] is numerically identical with the values of P_syst_ which are shown in the line “Arterial pressure”. It is evident that pulse pressure is not a fixed value when the resistance is changing; pulse pressure grows linearly with increment of growth equal to 1.25.

**Table 3.**
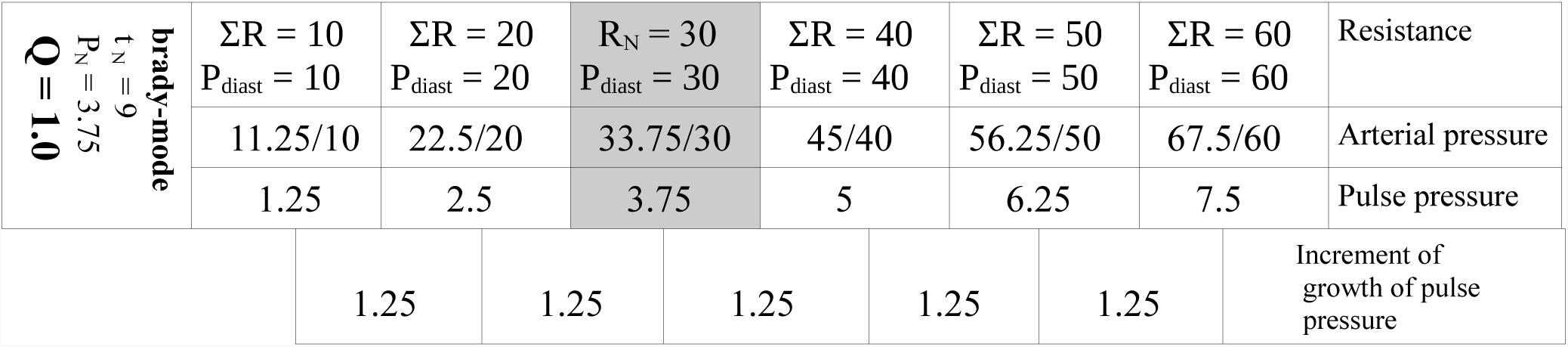
The steady rising of the resistance (which is identical to diastolic pressure when Q = 1.0) results in growth of pulse pressure; the represented brady-mode of circulation is shortly described by information in the upper left cell; the gray column indicates the state of brady-mode when circulation is not deviated by shift of the resistance (it is the starting point, i.e., the normal state for brady-mode).

As far as, - at the general case, - we do not know if the measured diastolic pressure is accompanied by Q = 1.0 or not, it is pertinent to investigate also the extreme modes of circulation (tachy-mode and brady-mode) with forced volume flow rate (Q = 1.2). We are going to find out – how the increment of growth of pulse pressure is influenced by the forcing of volume flow rate.

Tab.4, the main part, is based on data from Fig.4 [1]; the upper small table is the central part of Tab.2 and it is used for comparison (tachy-mode with volume flow rate not forced, i.e., Q = 1.0); the main table reflects the steps of reorganization of tachy-mode of circulation with forced volume flow rate (t_N_ =3, P_N_ =18, Q = 1.2) as a response upon the rising and falling of the resistance which now differs from P_diast_. Three steps of response of L (data from Fig.4 [1]; L is assumed numerically equal to P_syst_) upon the rising of the resistance and two steps of response of L upon the falling of the resistance are represented as values of systolic pressure (see the line “Arterial pressure”). Tab.4 shows the series of ΣR which now differs from the values of P_diast_ (both are represented in the line “Resistance”); the values of P_diast_ can also be found in the line “Arterial pressure”; the series of L (L18, L36, L54, L72, L90, L108) from Fig. 4 [1] is numerically identical with the values of P_syst_ which are shown in the line “Arterial pressure”. It is evident that pulse pressure is not a fixed value when the resistance (and diastolic pressure) are changing; pulse pressure grows linearly with increment of growth equal to 6. When comparing to the increment of growth of pulse pressure which persists at tachy-mode with Q = 1.0 (i.e., the quantity of 5 from Tab. 2), we may notice that new quantity of 6 (Tab.4) is higher; it means that pulse pressure is growing still linearly but the slope of the function is more steep.

**Table 4.**
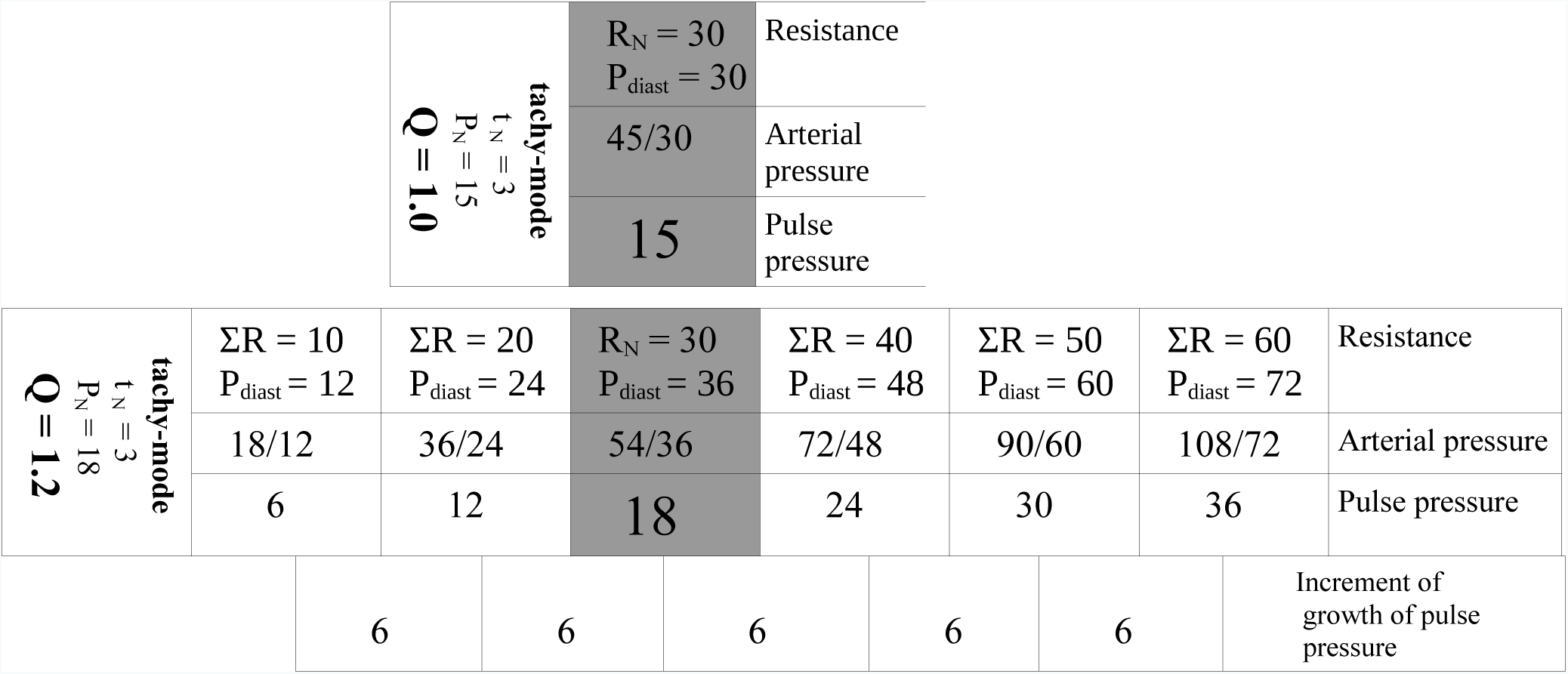
The steady rising of the resistance (which differs from diastolic pressure when Q = 1.2) results in growth of pulse pressure. The upper small table is the central part from the Tab.3 (for comparison); the main table represents tachy-mode of circulation with forced volume flow rate (basic parameters are given at the upper left cell); the gray column indicates the state of circulation when tachy-mode with Q = 1.2 is not deviated by shift of the resistance (it is the starting point, i.e., the normal state for tachy-mode with Q = 1.2).

Tab.5, the main part, is based on data from Fig.5 [1]; the upper small table is the central part of Tab.3 and it is used for comparison (brady-mode with volume flow rate not forced, i.e. Q = 1.0); the main table reflects the steps of reorganization of brady-mode of circulation with forced volume flow rate (t_N_ =9, P_N_ =4.5, Q = 1.2) as a response upon the rising and falling of the resistance which now differs from P_diast_. Three steps of response of L (data from Fig.5 [1]; L is assumed numerically equal to P_syst_) upon the rising of the resistance and two steps of response of L upon the falling of the resistance are represented as values of systolic pressure (see the line “Arterial pressure”). Tab.5 shows the series of ΣR which now differs from the values of P_diast_ (both are represented in the line “Resistance”); the values of P_diast_ can also be found in the line “Arterial pressure”; the series of L (L13.5, L27, L40.5, L54, L67.5, L81) from Fig. 5 [1] is numerically identical with the values of P_syst_ which are shown in the line “Arterial pressure”. It is evident that pulse pressure is not a fixed value when the resistance (and diastolic pressure) are changing; pulse pressure grows linearly with increment of growth equal to 1.5. When comparing to the increment of growth of pulse pressure which persists at brady-mode with Q = 1.0 (i.e., the quantity of 1.25 from Tab.3), we may notice that new quantity of 1.5 (Tab.5) is higher; it means that pulse pressure is growing still linearly but the slope of the function is more steep.

**Table 5.**
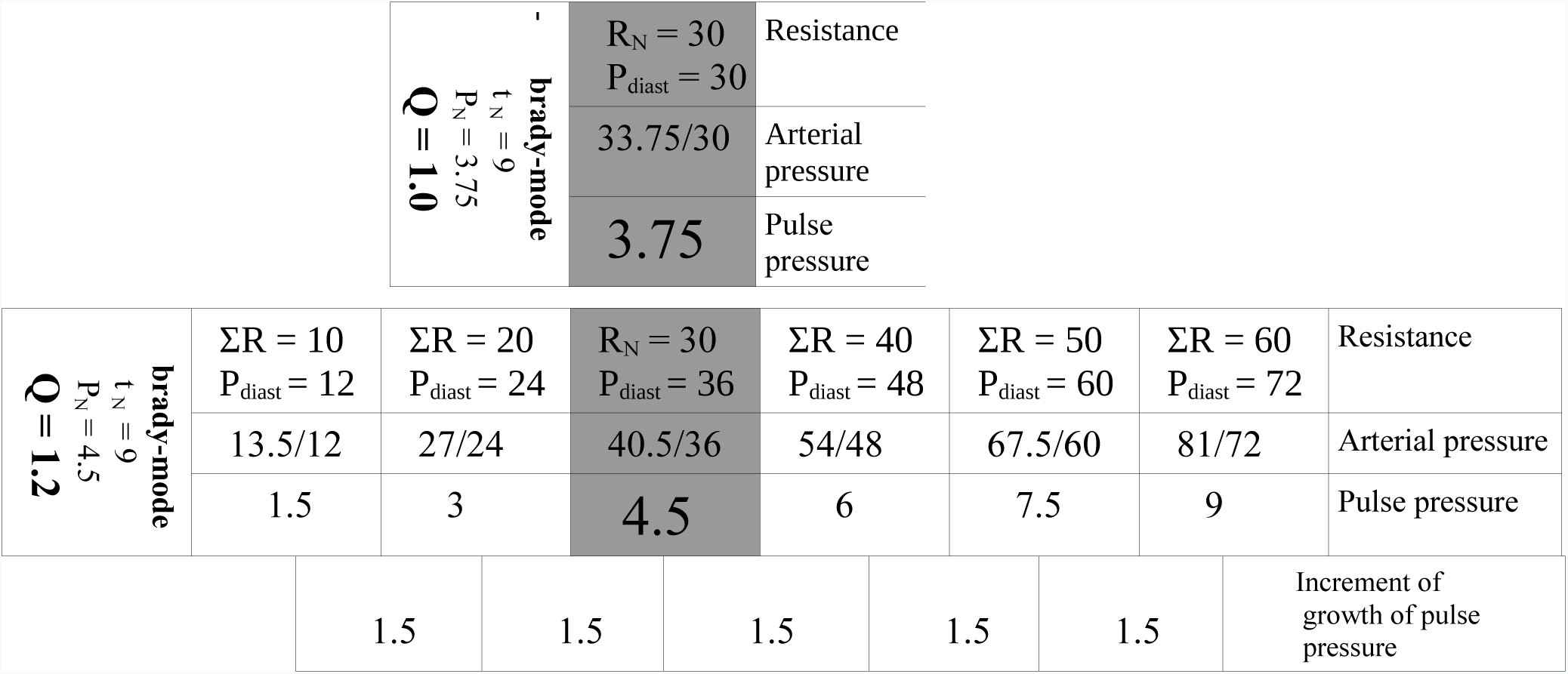
The steady rising of the resistance (which differs from diastolic pressure when Q = 1.2) results in growth of pulse pressure. The upper small table is the central part from the Tab.3 (for comparison); the main table represents brady-mode of circulation with forced volume flow rate (basic parameters are given at the upper left cell); the gray column indicates the state of circulation when brady-mode with Q = 1.2 is not deviated by shift of the resistance (it is the starting point, i.e., the normal state for brady-mode with Q = 1.2).

Therefore, five examples demonstrate the linear growth of pulse pressure while diastolic pressure is growing with steady increment. The dependency is based on calculation of end-diastolic deformation L (end-diastolic volume) which outstrips the steady elevation of the resistance (analysis of functions see below). The transition from L to systolic volume is the approximated function of direct proportionality (Frank-Starling law) and, consequently, the transition from systolic volume to systolic pressure is unknown only. The question is: either the latter transition duplicates the direct proportionality of Frank-Starling law (maybe with slightly different coefficient of proportionality) or the transition (from systolic volume to systolic pressure) distorts the above direct proportionality of Frank-Starling law? We have supposed that the transition is also a directly proportional function with coefficient of proportionality equal to1, i.e., we have numerically equalized systolic volume (and end-diastolic volume, respectively, as far as coefficient of proportionality of Frank-Starling law was assumed equal to 1) with systolic pressure. As a result, we see the series of ratios of systolic and diastolic pressure (arterial pressure) and series of values of pulse pressure which all demonstrate the one specific feature: the outstripping growth of systolic pressure comparing to elevation of diastolic pressure (the increment of growth of pulse pressure is steady and differs from zero).

The appearance of outstripping effect is evident when the calculation of the end-diastolic volume, L, is referred to the formula 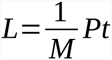, where M is a coefficient of dynamic viscosity (which is constant and equal to 1), t is the duration of diastole (which elongation is a response to elevation of resistance and which restoration to the normal value happens when the venous pressure rises, i.e., t is a constant value) and P is the venous pressure (which is the only variable that retains its value while being increased; the increment of P is steady) [1]. Therefore, the growth of L is accomplishing due to multiplication of P (the value is formed by addition of the steady increment to the previous value) by numerical constant; in that way, we have constructed the function of direct proportionality with substantial coefficient of proportionality which determines the certain angle of linear function relatively the abscissa axis. The growth of the resistance is accomplishing due to addition of the steady increment ΔR to the previous value of resistance and, consequently, the elevation of diastolic pressure which is represented by *P*_*diast*_ = *Q*·*ΣR* can be considered the function of direct proportionality with Q as a factor; this function has its own slope relatively the abscissa axis. It is obvious that the functions *L* = *f(P)* and *P*_*diast*_=*f*(*ΣR*) possess the individual angles relatively the abscissa axis and the discrepancy between the values of the functions (between L and P_diast_) is increasing while the arguments (P and ΣR) are changing due to their steady increments. The above discrepancy between end-diastolic volume L, - which we assume proportionate to systolic pressure with coefficient of proportionality equal to 1 (via proportionality to systolic volume with coefficient of proportionality equal to 1), - and diastolic pressure P_diast_ has the interpretation of pulse pressure, i.e., the growth of systolic pressure P_syst_ outstrips the growth of P_diast_ if the above substantial coefficient of the linearity which operates at *P*_*syst*_=*f*(*P*) prevails over Q at *P*_*dias*_=*Q · ΣR*. Certainly, the ascending limbs of two different parabolas, for instance, may demonstrate the analogous discrepancy but the steady increment of outstripping of values proves that both domains restrict the ranges of functions where these functions can be approximated to linearity. Besides, in such a case, the function *P*_*diast*_ = *Q*·*ΣR* pretends to be a non-linear function which resembles a square function but the growth of the volume flow rate in parallel with the resistance is hardly probable.

If we observe something of the kind in reality it means that calculations on the basis of the model [1] are true and the transition from systolic volume to systolic pressure is linear (or maybe circulation uses only quasi-linear portion of the function which generally is not linear just like in case of Frank-Starling law where linearity is only the approximation of an ascending limb of square function [2]). The most simple way is to draw the attention to clinical practice of measuring of arterial pressure. What combinations of systolic and diastolic pressures, - which we consider ordinary either at conditions of rest or at physical exertions, have we got used to observe? Tab.6 proposes the data which is trivial and approximate but the information reflects the trends which are undoubtedly observable, first of all, within sports medicine.

**Table 6.** The ordinary clinical gradation of arterial hypertension (no connection with renal pathology or valvular defects); quantified values are approximate but quite observable (either within essential arterial hypertension or within responses to physical exertion).

|  | Diastolic pressure | Increment of growth of diastolic pressure | Pulse pressure | Increment of growth of pulse pressure |
| --- | --- | --- | --- | --- |
| “Low arterial pressure”<br>90/60 | 60 |  | 30 |  |
| Normal arterial pressure<br>120/70 - 120/80 | 70-80 | 10-20 | 40-50 | 10-20 |
| Heightened arterial pressure<br>140/90 - 150/90 | 90 | 10-20 | 50-60 | 10 |
| High arterial pressure<br>160/100 - 200/110 | 100-110 | 10-20 | 60-90 | 10-30 |

The “low arterial pressure” is considered 90/60 (pulse pressure is equal to 30). The average normal arterial pressure is 120/80 or 120/70 (pulse pressure is 40-50), - i.e., the raise of diastolic pressure by 10–20 units results in elevation of pulse pressure by 10–20 units. The heightening of arterial pressure begins from 140/90, 150/90 and, consequently, the diastolic pressure raises by 10–20 comparing to normal scope (70–80); simultaneously pulse pressure reaches the values of 50–60, - i.e., it elevates from 40–50 up to 50–60 (with increment of growth equal to 10 units). The next step of heightening of arterial pressure is stipulated by the observing values of diastolic pressure within 100–110; the values of systolic pressure which accompany such level of diastolic pressure are within range of 160–200 (or higher). Therefore, standard raise of diastolic pressure by 10–20 units results in the increasing of pulse pressure up to 60–90 and it exceeds the previous values of 50–60 by 10–30 units, and it is quite close to the previous increments of growth of pulse pressure (which were equal to 10–20 and 10). Certainly, something happens at the upper extreme portion of functions that are responsible for transition: 1) from diastolic volume to systolic volume and 2) from systolic volume to systolic pressure, – i.e., some non-linearity appears when volumes and pressures exceed some operating range.

The above numerical argumentation obtained from such primitive source like approximate observations available to each physician can be considered forcible omitting statistics because the above numerical argumentation illustrates the phenomenon. The phenomenon does not estranges from statistical analysis but phenomenon permits to manage without it due to the evidence; at least, the phenomenon realizes the attempt to become explicable when it corresponds to some calculations and modeling. The coincidence of theoretical prognostication and practical data is just in favor of the suggested model of one-loop circulation (described in [1] and supplemented by the present work) which can be exploited for some more complex simulation.

